# The roles of distinct Ca^2+^ signaling mediated by Piezo and inositol triphosphate receptor (IP3R) in the remodeling of E-cadherin during cell dissemination

**DOI:** 10.1101/2021.11.10.467957

**Authors:** Alejandra J.H. Cabrera, Barry M. Gumbiner, Young V. Kwon

## Abstract

Given the role of E-cadherin (E-cad) in holding epithelial cells together, the inverse relationship between E-cad levels and cell invasion has been perceived as a principle underlying the invasiveness of tumor cells. In contrast, our study employing the *Drosophila* model of cell dissemination demonstrates that E-cad is necessary for the invasiveness of *Ras^v12^*-transformed cells *in vivo*. *Drosophila* E-cad/β-catenin disassembles at adherens junctions and assembles at invasive protrusions—the actin- and cortactin-rich invadopodia-like protrusions associated with breach of the extracellular matrix (ECM)—during cell dissemination. Loss of E-cad attenuates dissemination of *Ras^v12^*-transformed cells by impairing their ability to compromise the ECM. Strikingly, the remodeling of E-cad/β-catenin subcellular distribution is controlled by two discrete intracellular calcium signaling pathways: Ca^2+^ release from endoplasmic reticulum via the inositol triphosphate receptor (IP3R) disassembles E-cad at adherens junctions while Ca^2+^ entry via the mechanosensitive channel Piezo assembles E-cad at invasive protrusions. Thus, our study provides molecular insights into the unconventional role of E-cad in cell invasion during cell dissemination *in vivo* and describes the discrete roles of intracellular calcium signaling in the remodeling of E-cad/β-catenin subcellular localization.

## Introduction

E-cadherin (E-cad), the main component of adherens junctions, helps to hold epithelial cells together (1–5). To invade and migrate, cells disintegrate adherens junctions to free themselves from neighboring cells (3, 6, 7). When epithelial cells invade and migrate during development, they undergo epithelial mesenchymal transition (EMT), a process that allows them to acquire the properties of mesenchymal cells (8, 9). Consistent with the role of E-cad, loss of E-cad is considered to be a hallmark of EMT (10, 11). Frixen et al. showed that loss of E-cad is also important for the ability of carcinoma cells to invade and migrate in culture (12). These observations coincide with clinical reports, indicating that E-cad is frequently reduced in epithelial cell cancers (13, 14). Therefore, reduction of E-cad is thought to promote cancer cell invasion and metastasis (12, 15–22). However, a number of clinical studies have reported that E-cad is expressed in multiple metastatic tumors (23–26). Notably, Padmanaban et al. have recently demonstrated that E-cad is required for metastasis in multiple invasive ductal carcinoma models by helping cancer cells to survive after dissemination (15), suggesting that E-cad might play a more complicated, not fully defined, role in metastasis of cancer cells.

We attempted to address the role of E-cad in the invasiveness of transformed cells in a native microenvironment by employing our *Drosophila* model of *Ras^V12^-* transformed cell dissemination (27). Expression of *Ras^V12^* in adult intestinal stem cells (ISCs) and enteroblasts (EBs) using the conditional GAL4 driver *esg-GAL4*, *UAS-GFP*, *tub-GAL80^ts^* (*esg^ts^*; see methods) makes *Ras^V12^*-expressing intestinal epithelial cells (*Ras^V12^* cells) disseminate basally from the midgut and enter the hemocoel—the primary cavity containing circulatory fluid (27, 28) (also see Figure for reviewer). Note that *Ras^V12^* cells also delaminate apically toward lumen and then are presumably excreted (27). During dissemination, *Ras^V12^* cells generate actin- and cortactin-rich invasive protrusions functionally and structurally reminiscent of invadopodia observed in cancer cells to breach the ECM and the visceral muscle (VM) layer (27, 29–32). After compromising the integrity of the ECM and the VM, *Ras^V12^* cells can then migrate into the circulation by bleb-driven movement. Depletion of *cortactin* or the mechanosensitive channel *piezo* specifically in *Ras^V12^* cells attenuates dissemination of *Ras^V12^* cells by impairing their ability to invade and migrate across the ECM and the VM. Thus, this model allows us not only to observe the cell dissemination process in a native context at the cellular and molecular levels, but also to test the function of a gene in the cell dissemination process using the advanced genetic tools available in *Drosophila*.

Our findings demonstrate that E-cad is necessary for the invasiveness of *Ras^V12^* cells during cell dissemination *in vivo*. Our observations suggest that subcellular E-cad is redistributed in disseminating *Ras^V12^* cells: E-cad disassembles at adherens junctions–a process that recapitulates the functional consequence of E-cad loss during EMT–and assembles at invasive protrusions, which is not a conventional location for E-cad. We show that two distinct intracellular calcium signaling pathways mediated by the inositol triphosphate receptor (IP3R) and Piezo control disassembly of E-cad at adherens junctions and assembly at invasive protrusions, respectively. Therefore, our study provides new insights into the role of E-cad in invasiveness and elucidates the mechanisms by which two intracellular calcium signaling pathways differentially control the disassembly and assembly of E-cad to aid cell dissemination.

## Results

### E-cadherin is required for dissemination of *Ras^V12^* cells

Since our knowledge of the role of E-cad in cell dissemination is limited, we embarked our study by testing whether E-cad is necessary using the *Drosophila Ras^V12^-* induced cell dissemination model (Figure for reviewer) (27). At day 2 of *Ras^V12^* expression, disseminated *Ras^V12^* cells residing at the outer surface of the posterior midguts can be quantified to assess cell dissemination (27). Depletion of *Drosophila E-cadherin (DE-cad*; also known as *shotgun* [*shg*]) in *Ras^V12^* cells by expressing *DE-cad* RNA interference (RNAi) (*JF02769* and *HMS00693*) with *esg^ts^* didn’t dramatically alter the overall distribution of *Ras^V12^* cells in the posterior midguts (Figure S1a). However, *E-cad* depletion in *Ras^V12^* cells almost completely eliminated disseminated *Ras^V12^* cells detected at the outer surface of the posterior midguts (Figure 1a and b). Given the inverse relationship between E-cad levels and cell migration in cultured cancer cells (12), we also increased DE-cad levels in *Ras^V12^* cells by expressing *UAS-DE-cad* with *esg^ts^*. E-cad overexpression only partially reduced dissemination of *Ras^V12^* cells (Figure 1b). At day 2 of *Ras^V12^* expression, *Ras^V12^* cells also apically delaminate (Figure for reviewer). Interestingly, *DE-cad* depletion didn’t impair apical delamination of *Ras^V12^* cells (Figure S1b), suggesting that inability to move out from the epithelium was not likely to account for the cell dissemination defect caused by *DE-cad* depletion in *Ras^V12^* cells. Altogether, we concluded that E-cad was necessary for cell dissemination.

**Figure 1.**
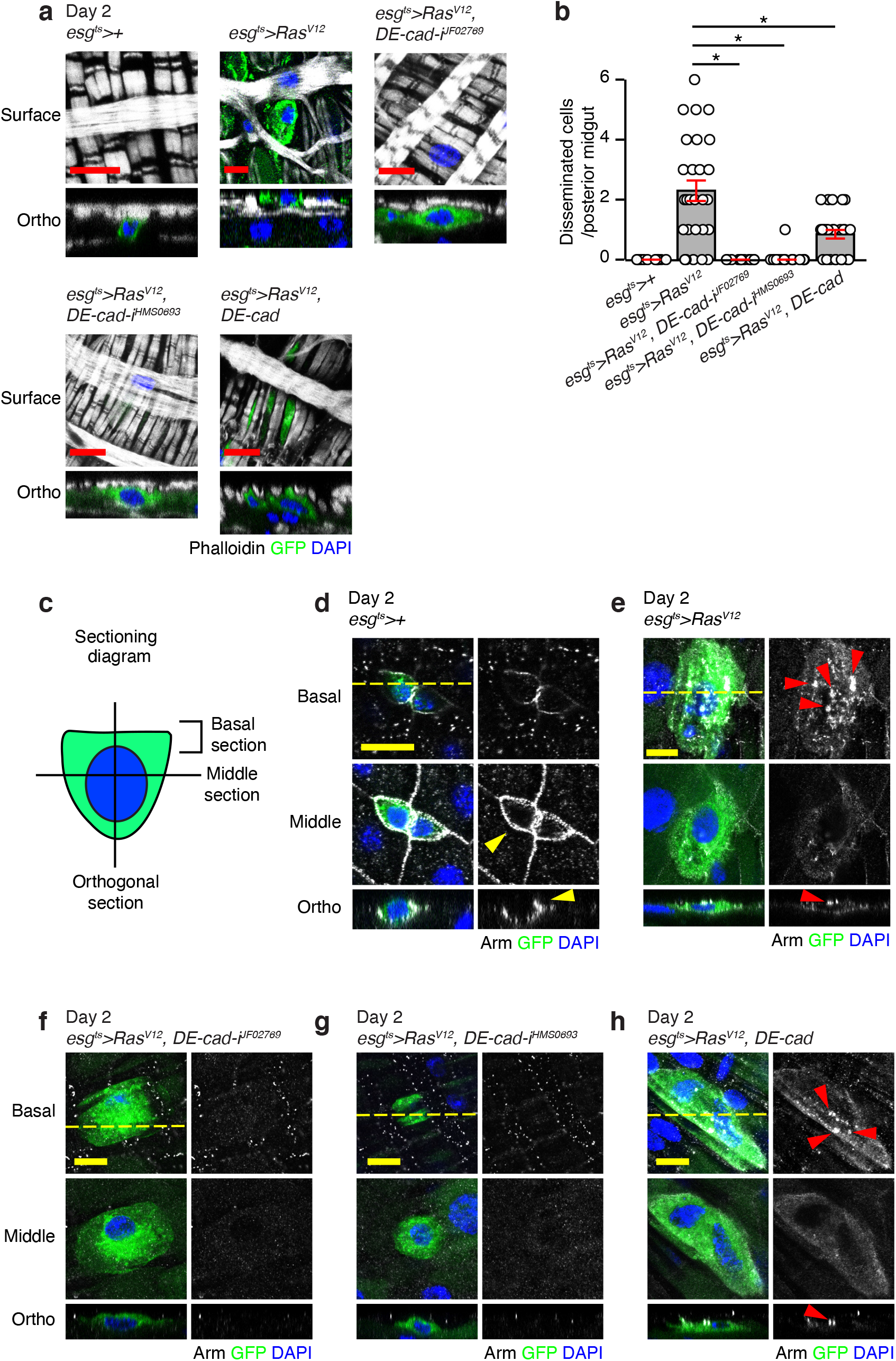
DE-cadherin is necessary for cell dissemination. **a,** Surface views and orthogonal images of representative cells; *esg^ts^>+*, control cell residing in the intestinal epithelium; *esg^ts^>Ras^V12^*, representative disseminated *Ras^V12^* cell residing at the outer surface of the visceral muscle; *esg^ts^>Ras^V12^*, *DE-cad-i^JF02769^* and *esg^ts^>Ras^V12^*, *DE-cad-i^JF02769^* cell, representative DE-cad depleted *Ras^V12^* cells residing in the intestinal epithelium; *esg^ts^>Ras^V12^*, *DE-cad*, representative *Ras^V12^* cells overexpressing *DE-cad* in the intestinal epithelium. The visceral muscle is visualized with Phalloidin (gray), *ras^V12^* cells are marked with GFP (green), and nuclei are stained with DAPI (blue). Scale bar, 10 μm. **b,** Quantification of disseminated cells residing on the outer VM surface at the posterior midguts. From left to right, *P*=4.3971 x 10^-9^ (*Ras^V12^* - *Ras^V12^*, *DE-cad-i^JF02769^*), *P*=1.0966 x 10-^9^ (*Ras^V12^* - *Ras^V12^*, *DE-cad-i^HMS0693^*), *P*= 5.9459 x 10^-6^ (*Ras^V12^* - *Ras^V12^*, *DE-cad*). *n*=12 (*esg^ts^*), *n*=27 (*esg^ts^>Ras^V12^*), *n*=13 (*esg^ts^>Ras^V12^*, *DE-cad-i^JF02769^*), *n*=16 (*esg^ts^>Ras^V12^*, *DE-cad-i^HMS0693^*), *n*=26 (*esg^ts^>Ras^V12^*, *DE-cad*) biological replicates. Mean±SEM are shown with individual data points. Statistical analysis was performed using one-way ANOVA with post-hoc Tukey HSD. Asterisks indicate statistical significance (\**P*<0.05). **c-h**, Representative images of cells stained with anti-Arm antibody (gray). Genotypes are shown. Top images representing the basal section of the cells are projections of 2-4 very basal z-stacks. Middle images show the representative cross sections capturing adherens junctions of the cells. In the orthogonal views, the basal side of cells is positioned upward. Red arrowheads indicate Arm signals detected at cell-cell junctions, and yellow arrowheads indicate Arm signals detected as distinct puncta at the basal side of *Ras^V12^* cells. Scale bar, 10 μm.

### The subcellular distribution of DE-cad/Arm is remodeled during *in vivo* cell dissemination

To gain insight into how adherens junctions were affected during cell dissemination, we scrutinized the subcellular distribution of DE-cad in *Ras^V12^* cells. At day 1 of *Ras^V12^* expression, most of the *Ras^V12^* cells stay in the midgut epithelium, resulting in hyperplasia (27). At day 2, *Ras^V12^* cells basally disseminate from the midgut while a significant number of *Ras^V12^* cells also delaminate apically toward the lumen (27) (Figure S1b and Figure for reviewer). When we stained control and *Ras^V12^* cells at day 1 with anti-DE-cad antibody, strong signals were detected at the lateral side (Figure S2a), which was recapitulated by staining for the *Drosophila* β-catenin ortholog Armadillo (Arm) (Figure S2b). Since Arm staining yielded much less background signals at the VM (Figure S2a and b, nonspecific signals in the VM layer is indicated with asterisks), we decided to use Arm staining as the primary method to assess E-cad subcellular distribution. At day 2 of *Ras^V12^* expression, Arm signals were no longer clearly visible at the lateral side of *Ras^V12^* cells while cytoplasmic Arm signals were increased (Figure 1e, middle section), suggesting that disassembly of adherens junctions rather than reduction in E-cad levels was a mechanism to aid dissemination of *Ras^V12^* cells. Notably, we detected Arm signals as discrete puncta at the basal side of *Ras^V12^* cells (Figure 1e, basal section, red arrowheads). These basal Arm signals were not visible in control cells (Figure 1d) or *Ras^V12^* cells at day 1 (Figure S2b). E-cad was also detected as puncta at the basal side of *Ras^V12^* cells when we stained with anti-E-cad antibody or used the DE-cad protein trap line (*shg^mTomato^*) producing DE-cad-mTomato under the control of its own promotor (Figure S2c and d). Additionally, Arm colocalized with DE-cad-mRFP at the basal side of *Ras^V12^* cells (Figure S2d), and *DE-cad* depletion impaired formation of Arm puncta at the basal side of *Ras^V12^* cells (Figure 1f and g, middle sections), indicating that the formation of these puncta was dependent on E-cad. Overexpression of DE-cad didn’t stabilize adherens junctions nor impair formation of Arm puncta at the basal side of *Ras^V12^* cells (Figure 1h, basal and middle sections). Taken together, these observations demonstrate that DE-cad/Arm undergoes extensive remodeling during dissemination of *Ras^V12^* cells. DE-cad disassembles at adherens junctions and assembles as puncta at the basal side of *Ras^V12^* cells, which doesn’t appear to be simply controlled by DE-cad levels.

### DE-cad/Arm assembles at invasive protrusions and is required for the invasiveness of *Ras^V12^* cells

The unconventional DE-cad/Arm puncta at the basal side of *Ras^V12^* cells are reminiscent of *Drosophila* invasive protrusions, which are actin- and cortactin-rich protrusions associated with degradation of the ECM and the VM (27). Thus, we tested whether DE-cad/Arm was localized at invasive protrusions. Considering the possible limitations associated with various actin markers (33, 34), we decided to use three different actin reporters, Lifeact-mRFP (34, 35), mCherry-Moe.ABD (36), and actin-mRFP (37) to visualize invasive protrusions (Figure 2a, arrowheads). Importantly, Arm signals co-localized to the puncta marked by the actin markers (Figure 2a, arrowheads). Thus, these results indicate that DE-cad/Arm is a new component of invasive protrusions.

**Figure 2.**
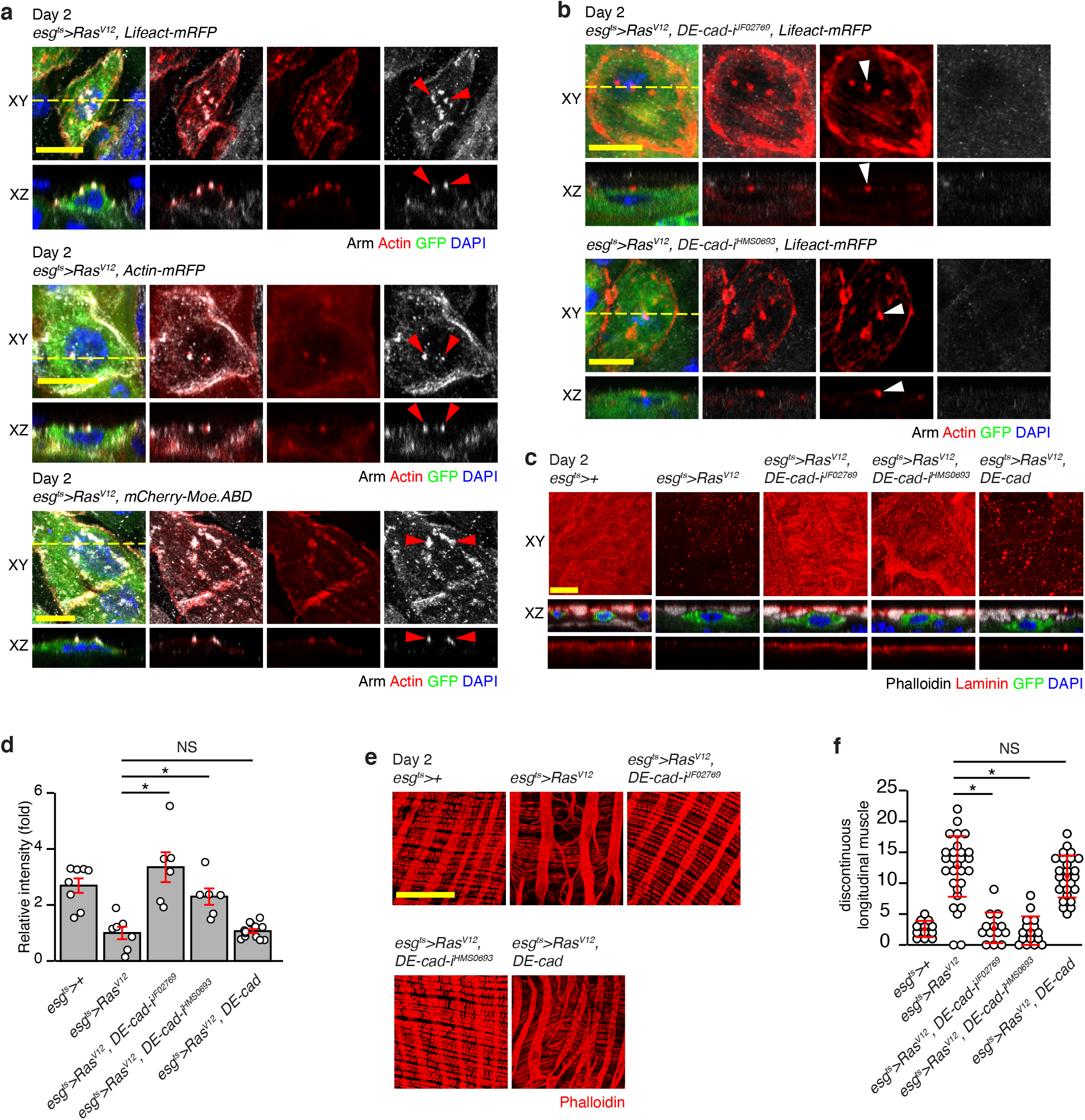
DE-cad/Arm assembles at invasive protrusions and controls their function. **a,** Basal sections (xy) and orthogonal views (xz) of *Ras^V12^* cells. Transgenes are induced for 2 days. Actin (red) is visualized *UAS-Lifeact-mRFP, UAS-Actin-mRFP*, and *UAS-mCherry-Moe.ABD*. Arm (gray) was detected by staining with anti-Arm antibody. In the orthogonal views, the basal side of cells is positioned upward. Co-localization of an actin marker and Arm is indicated with arrowheads. Nuclei are stained with DAPI (blue). Scale bars, 10 μm. **b,** Lifeact-mRFP signals (red) in *Ras^V12^*, *DE-cad RNAi* cells. Lifeact-mRFP puncta located at the basal side of *Ras^V12^* cells represent invasive protrusions (arrowheads). Two DE-cad RNAi lines (*JF02769* and *HMS0693*) are used to knockdown *DE-cad* in *Ras^V12^* cells. Top images (xy), basal sections; bottom images (xz), orthogonal sections. **c,** Laminin B1 staining. Top images, top views (xy); bottom images, orthogonal sections (xz). *Ras^V12^* cells are marked with GFP (green). All genotypes were stained with anti-Laminin B1 antibody (red), Phalloidin (gray) and DAPI (blue). Scale bars, 10 μm. **d,** Quantification of Laminin B1 signals. From left to right, *P*=8.9517 x 10^-06^ (*Ras^V12^* - *Ras^V12^*, *DE-cad-i^JF02769^*), *P*=0.0186 (*Ras^V12^* - *Ras^V12^*, *DE-cad-i^HMS0693^*), *P*=0.9996 (*Ras^V12^* - *Ras^V12^*, *DE-cad*). *n*=8 (*esg^ts^*), *n*=7(*esg^ts^*>*Ras^V12^*), *n*=6 (*esg^ts^*>*Ras^V12^*, *DE-cad-i^JF02769^*), *n*=6 (*esg^ts^*>*Ras^V12^*, *DE-cad-i^HMS0693^*), *n*=13 (*esg^ts^*>*Ras^V12^*, *DE-cad*) biological replicates. **e,** Visceral muscle (VM) at the posterior midguts. VM (red) is visualized with phalloidin. Scale bars, 50 μm. **f,** Quantification of longitudinal muscle breaks. From left to right, *P*=5.4445 x 10^-13^ (*Ras^V12^* - *Ras^V12^*, *DE-cad-i^JF02769^*), *P*=9.2371 x 10^-14^ (*Ras^V12^* - *Ras^V12^*, *DE-cad-i^HMS0693^*), *P*=0.48779 (*Ras^V12^* - *Ras^V12^*, *DE-cad*). *n*=12 (*esg^ts^*), *n*=27(*esg^ts^*>*Ras^V12^*), *n*=13 (*esg^ts^>Ras^V12^*, *DE-cad-i^JF02769^*), *n*=16 (*esg^ts^>Ras^V12^*, *DE-cad-i^HMS0693^*), *n*=26 (*esg^ts^>Ras^V12^*, *DE-cad*) biological replicates. In **d** and **f**, mean±SEMs are shown with individual data points. Data were analyzed by using one-way ANOVA with post-hoc Tukey HSD. Asterisks indicate statistical significance (*P*<0.05). Transgenes were expressed with *esg^ts^* for 2 days by shifting to 29°C.

We then tested whether DE-cad was required for formation of invasive protrusions. Expression of *DE-cad* RNAi with *esg^ts^* did not alter the formation of Lifeact-RFP puncta at the basal side of *Ras^V12^* cells (Figure 2b), indicating that DE-cad is not crucial for the formation of invasive protrusions. Notably, *DE-cad* depletion impaired *Ras^V12^* cells’ ability to degrade the ECM (Figure 2c and d). At day 2 of *Ras^V12^* expression, the laminin layer was almost completely degraded (Figure 2c and d). In contrast, the laminin layer remained intact when *DE-cad* was depleted in *Ras^V12^* cells (Figure 2c and d). *Drosophila* invasive protrusions have been shown to damage the VM layer–manifested by occasional breakages of the longitudinal muscles that normally span the whole posterior part of the midgut (27) (Figure 2e and f). *DE-cad* depletion in *Ras^V12^* cells significantly reduced the longitudinal muscle breakages (Figure 2e and f). Note that DE-cad overexpression in *Ras^V12^* cells couldn’t suppress the laminin degradation or the longitudinal muscle breakage phenotypes (Figure 2c-f), indicating that an increase in DE-cad levels was not sufficient to impair the function of invasive protrusions. These results show that DE-cad plays an essential role in the invasiveness of *Ras^V12^* cells by controlling the function of invasive protrusions. Without DE-cad, *Ras^V12^* cells lose their ability to compromise the ECM and the VM for dissemination.

### Intracellular calcium signaling via the PLC-IP3R-CAMK pathway is required for disassembly of DE-cad/Arm at adherens junctions in *Ras^V12^* cells

Since our findings suggest that the subcellular distribution of DE-cad/Arm is actively remodeled in disseminating cells, we sought to elucidate the signaling mechanisms underlying the remodeling process. Our previous report demonstrated that calcium signaling mediated by the mechanosensitive cation channel Piezo was essential for dissemination of *Ras^V12^* cells (27), which led us to investigate the importance of intracellular calcium signaling in cell dissemination. IP3R increases cytosolic calcium levels by releasing calcium from the endoplasmic reticulum (ER) in response to its ligand inositol triphosphate (IP3), which is generated by phospholipase C (PLC)-mediated cleavage of phosphatidylinositol 4,5-bisphosphate (PIP2) (38, 39). At day 2 of *Ras^V12^* expression, a significant portion of *Ras^V12^* cells basally disseminate and apically delaminate (27). Therefore, *Ras^V12^* cells form tumors at low frequency at day 2 (Figure 3a and Figure S3a). Interestingly, *IP3R* depletion made *Ras^V12^* cells stay in the midguts, resulting in an increase in tumor formation at day 2 (Figure 3a and Figure S3a). Moreover, *IP3R* depletion attenuated dissemination (Figure 3b) and delamination of *Ras^V12^* cells (Figure S3b). These observations led us to speculate that intracellular calcium signaling mediated by IP3R might be important for disassembly of DE-cad/Arm at adherens junctions. Thus, we attempted to identify the pathway components using tumor growth as a readout. To identify the PLC functioning upstream of IP3R, we tested all three PLCs in the genome, and found that depletion of the PLCγ *small wing* (*sl*) in *Ras^V12^* cells led to tumor formation (Figure 3a and Figure S3c). Additionally, depletion of both *Ca^2+^/calmodulin*-*dependent protein kinase* (*CaMK*) *I* and *II* in *Ras^V12^* cells induced tumors (Figure 3a and Figure S3a). Knockdown of individual CaMKs in *Ras^V12^* cells induced tumors at a low frequency (Figure S3a), suggesting a redundancy in their functions. Notably, depletion of either *sl* or *CaMKI* and *II* also significantly suppressed dissemination of *Ras^V12^* cells (Figure 3b). These phenotypic similarities suggest that Sl, IP3R, and CaMKs might form a pathway to operate calcium release from the ER in *Ras^V12^* cells, which is crucial for dissemination of *Ras^V12^* cells.

**Figure 3.**
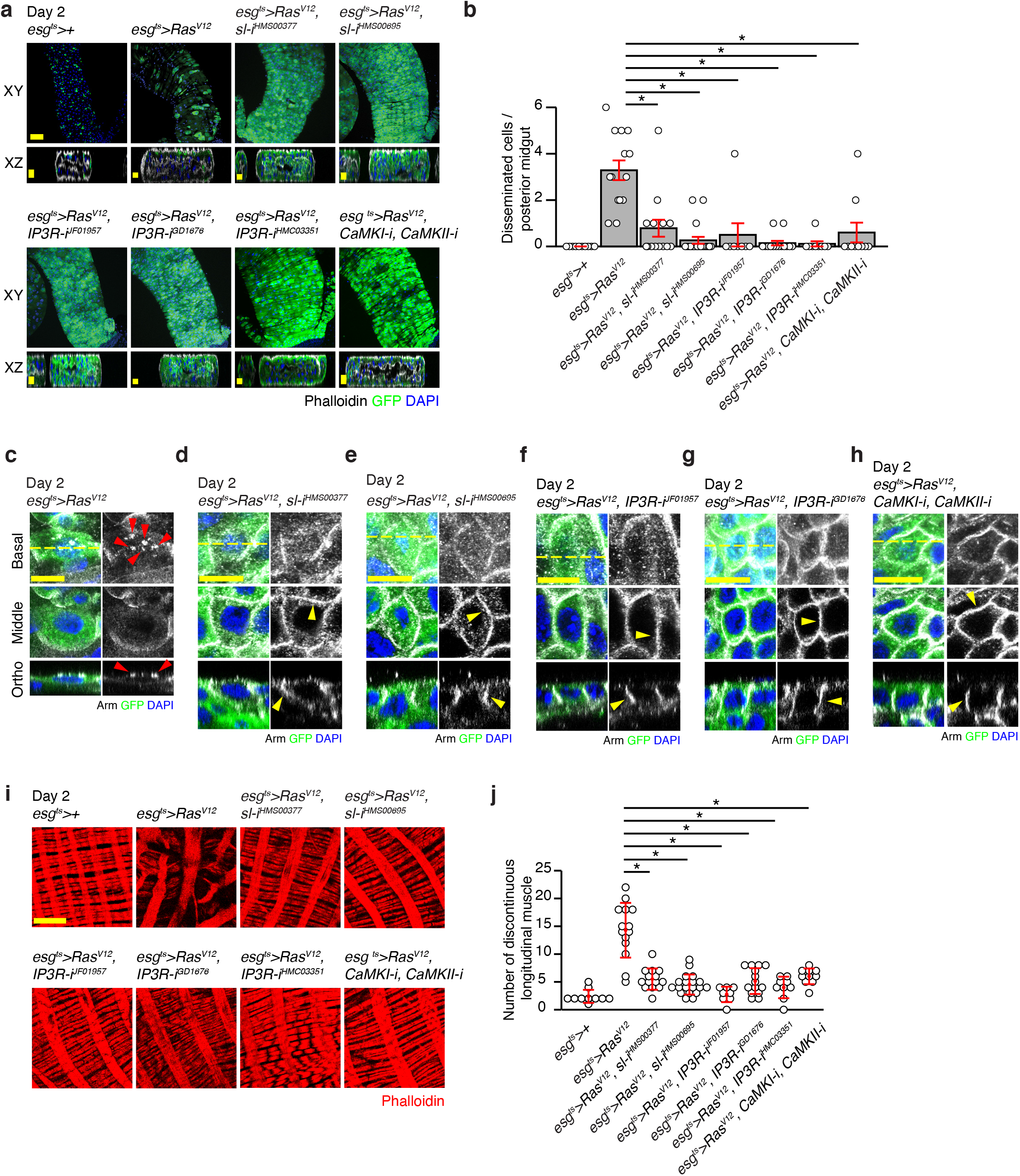
The PLC-IP3R-CAMK pathway is required for DE-cad/Arm disassembly from adherens junctions and assembly at invasive protrusions. **a,** Representative images of the posterior midgut showing tumor morphologies at day 2 of transgene expression with *esg^ts^*. *Ras^V12^* cells are marked with GFP (green), and nuclei are stained with DAPI (blue). VM is visualized with phalloidin (gray). Bottom panels are representative orthogonal views of the midguts. Scale bars, x: 50 μm and y: 10 μm. **b,** Quantification of disseminated cells. From left to right, *P*=3.0567 x 10^-10^ (*Ras^V12^* - *Ras^V12^*, *sl-i^HMS00377^*), *P*=7.3386 x 10^-14^ (*Ras^V12^* - *Ras^V12^*, *sl-i^HMS00695^*), *P*=2.2458 x 10^-09^ (*Ras^V12^* - *Ras^V12^*, *IP3R-i^JF01957^*), *P*=8.1712 x 10^-14^ (*Ras^V12^* - *Ras^V12^*, *IP3R-i^GD1676^*), *P*=1.9582 x 10^-12^ (*Ras^V12^* - *Ras^V12^*, *IP3R-i^HMC03351^*), *P*=6.6618 x 10^-10^ (*Ras^V12^* - *Ras^V12^*, *CaMKI-i*, *CaMKII-i*). **c-h,** Arm staining. All genotypes were stained with anti-Arm antibody (gray) and DAPI (blue). *Ras^V12^* cells are marked with GFP (green). Red arrowheads indicate basal Arm signal puncta, and yellow arrowheads show Arm signals at cell-cell junctions. Scale bar, 10 μm. **i**. VM detected by phalloidin staining (red). Scale bar, 25 μm. **J,** Quantification of longitudinal muscle breaks. From left to right, *P*=1.1258 x 10^-13^ (*Ras^V12^* - *Ras^V12^*, *sl-i^HMS00377^*), *P*=2.7534 x 10^-14^ (*Ras^V12^* - *Ras^V12^*, *sl-i^HMS00695^*), *P*=5.9730 x 10^-14^ (*Ras^V12^* - *Ras^V12^*, *IP3R-i^JF01957^*), *P*=7.8271 x 10^-14^ (*Ras^V12^* - *Ras^V12^*, *IP3R-i^GD1676^*), *P*=7.6605 x 10^-14^ (*Ras^V12^* - *Ras^V12^*, *IP3R-i^HMC03351^*), *P*=4.3077 x 10^-11^ (*Ras^V12^* - *Ras^V12^*, *CaMKI-i*, *CaMKII-i*). In **b** and **j**, *n*=10 (*esg^ts^*), *n*=14 (*esg^ts^*>*Ras^V12^*), *n*=14 (*esg^ts^*>*Ras^V12^*, *sl-i^HMS00377^*), *n*=19 (*esg^ts^*>*Ras^V12^*, *sl-i^HMS00695^*), n=8 (*esg^ts^*>*Ras^V12^*, *IP3R*-*i^JF01957^*), *n*=14 (*esg^ts^*>*Ras^V12^*, *IP3R-i^GD1676^*), *n*=9 (*esg^ts^*>*Ras^V12^*, *IP3R-i^HMC03351^*), *n*=10 (*esg^ts^*>*Ras^V12^*, *CaMKI-i*, *CaMKII-i*) biological replicates. Mean±SEMs are shown with individual data points. Statistical analysis was performed using one-way ANOVA with post-hoc Tukey HSD. Asterisks indicate statistical significance (*P<0.05). In all panels, transgenes are induced with *esg^ts^* for 2 days at 29°C.

To address whether the PLC-IP3R-CAMK pathway controls disassembly of DE-cad/Arm at adherens junctions, we assessed the subcellular distribution of Arm in *Ras^V12^* cells after depleting each component. In *Ras^V12^* cells, Arm signals disappeared from the lateral side (Figure 3c). In contrast, strong Arm signals were present at the lateral side of *Ras^V12^* cells when the PLC-IP3R-CAMK pathway was disrupted (Figure 3d-h, yellow arrowheads). Arm puncta were almost undetectable at the basal side of *Ras^V12^* cells when either *sl* or *IP3R* was depleted (Figure 3d-g). When both *CaMKs* were depleted, basal Arm puncta were only occasionally detected in some *Ras^V12^* cells (Figure 3h and Figure S4). Loss of DE-cad/Arm at invasive protrusions could result in an impairment of invasive protrusion’s function. Indeed, depletion of *sl*, *IP3R*, or *CaMKs* significantly impaired the *Ras^V12^* cells’ ability to disrupt the VM (Figure 3i and j). Thus, these observations suggest that the PLC-IP3R-CAMK pathway controls the remodeling of DE-cad/Arm subcellular distribution in *Ras^V12^* cells by inducing disassembly of DE-cad/Arm at adherens junctions. Additionally, the loss-of-function phenotypes indicate that the PLC-IP3R-CAMK pathway is also required for localization of DE-cad/Arm to invasive protrusions.

### The Piezo-calpain pathway controls the assembly of DE-cad/Arm at invasive protrusions

Piezo channels can initiate intracellular calcium signaling by transporting Ca^2+^ across the plasma membrane upon sensing mechanical cues in the microenvironment (40–43). Previous studies have elucidated the roles of Piezo channels in the invasiveness of transformed cells (27, 44, 45). In particular, Piezo channels has been shown to control matrix degradation, which is associated with invadopodia function (27, 45). Nevertheless, the molecular mechanism by which Piezo controls the invasiveness of transformed cells has not been elucidated. Since Piezo activation could increase cytosolic Ca^2+^, we asked whether depletion of *piezo* in *Ras^V12^* cells affects the redistribution of DE-cad/Arm. Interestingly, when *piezo* was depleted in *Ras^V12^* cells, discrete Arm signals were not detected at the cell boundary, and most of Arm signals were detected in the cytosol (Figure 4a). The absence of basal Arm puncta in *Ras^V12^*, *piezo RNAi* cells indicates that Piezo is required for assembly of DE-cad/Arm at invasive protrusions. Our previous study showed that the calcium-dependent non-lysosomal cysteine proteases, calpains, worked with Piezo to control the function of invasive protrusion and increase matrix metalloproteinase 1 (Mmp1) expression in *Ras^V12^* cells (27, 40). Depletion of *Calpain-A* (*CalpA*) was also sufficient to recapitulate the Arm subcellular distribution observed in *Ras^V12^*, *piezo RNAi* cells (Figure 4a), suggesting that calpains might transduce the calcium signaling induced by Piezo activation. The absence of Arm signals at adherens junctions could be caused by an impairment in either assembly of DE-cad/Arm or maintenance of existing DE-cad/Arm. To gain further insight, we assessed the effect of *piezo* depletion on adherens junctions at day 1 of *Ras^V12^* expression when adherens junctions were not disassembled (Figure 4b and Figure S2b). Interestingly, Arm signals were not clearly visible at the lateral side of *Ras^V12^* cells even at day 1 when *piezo* was depleted (Figure 4b). These observations support that an impairment in assembly of DE-cad/Arm at adherens junctions is likely to be responsible for the absence of Arm signals at the lateral side of *Ras^V12^*, *piezo RNAi* cells. Altogether, our results demonstrate that the Piezo-calpain pathway is required for assembly of DE-cad/Arm at invasive protrusions, implying that calcium signaling mediated by the Piezo-calpain pathway plays a distinct role in the DE-cad/Arm remodeling process. Considering the essential role of DE-cad in the function of invasive protrusions (Figure 2), we propose that controlling the assembly of DE-cad/Arm at invasive protrusions is the molecular mechanism by which Piezo controls the invasiveness of *Ras^V12^* cells.

**Figure 4.**
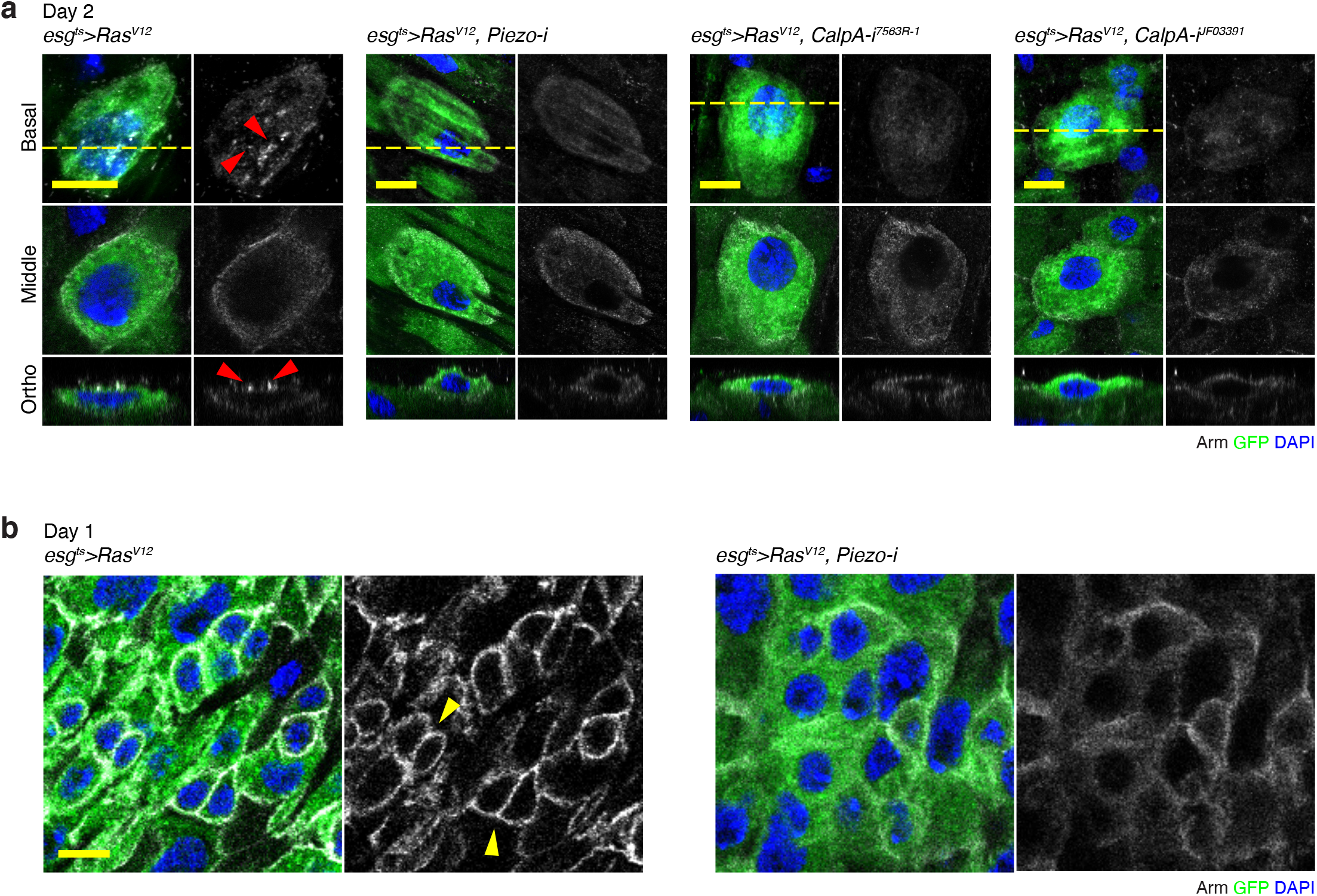
The Piezo-calpain pathway is required for assembly of DE-cad/Arm at invasive protrusions. **a,** Representative images of the basal, middle, and orthogonal sections of cells stained with anti-Arm antibody (gray). Arrowheads indicate Arm signals at invasive protrusions. The Piezo RNAi line (8486R-3) is also used in our previous study (27), and two RNAi lines (7563R-1 and JF03391) are used to knockdown *CalpA*. Transgenes were induced for 2 days with *esg^ts^*. Nuclei are stained with DAPI (blue). Scale bar, 10 μm. **b,** Arm staining (gray) of the midguts at day 1 of transgene expression with *esg^ts^*. DAPI (blue) marks nuclei. Junctions are indicated by arrowhead (yellow). Scale bar, 10 μm.

### An increase in cytosolic Ca^2+^ induces disassembly of DE-cad/Arm at adherens junctions

Our observations suggest that the two calcium signaling pathways–the PLC-IP3R-CAMK pathway and the Piezo-calpain pathway–elicit distinct effects on DE-cad/Arm redistribution in disseminating cells. To gain further insights into the roles of the calcium signaling pathways, we decided to assess the acute responses to manipulation of calcium signaling *ex vivo*. At day 1 of *Ras^V12^* expression, DE-cad/Arm is not disassembled from adherens junctions or assembled at invasive protrusions (Figure S2b). Thus, we attempted to assess how pharmacological manipulation of calcium signaling affects the Arm localization in *Ras^V12^* cells at day 1 of *Ras^V12^* expression. Sarco/endoplasmic reticulum Ca^2+^-ATPase (SERCA) transports Ca^2+^ from the cytosol to the ER, which keeps cytosolic Ca^2+^ concentration low (46). Thus, inhibition of SERCA leads to an increase in cytosolic Ca^2+^ and a depletion of Ca^2+^ in the ER. Thus, we used the inhibitor of SERCA, thapsigargin (TG), to induce a global increase in cytosolic Ca^2+^ (47). To inhibit Piezo-mediated calcium entry, we used gadolinium chloride (GdCl_3_), which inhibits the mechanically activated cationic currents (41). When we incubated day 1 *Ras^V12^* midguts in control media, Arm signals at adherens junctions were well-preserved (Figure 5a-c, H_2_O and DMSO). Strikingly, Arm signals were no longer concentrated at the lateral side of *Ras^V12^* cells when *Ras^V12^* midguts were treated with TG for 30 minutes (Figure 5b and c), indicating that an increase in cytosolic Ca^2+^ is sufficient to induce disassembly of DE-cad/Arm from adherens junctions in *Ras^V12^* cells. Although *piezo* depletion caused a loss of Arm signals from the lateral side of *Ras^V12^* cells *in vivo* (Figure 4b), transient GdCl_3_ treatment didn’t yield any discernable changes in Arm signals at the lateral side of *Ras^V12^* cells (Figure 5b and c), suggesting that inhibition of Piezo doesn’t delocalize DE-cad/Arm already existing at adherens junctions. Addition of GdCl_3_ didn’t significantly inhibit the TG-induced delocalization of Arm from the lateral side of *Ras^V12^* cells (Figure 5b and c). Thus, Piezo does not contribute to the TG-induced disassembly of DE-cad/Arm at adherens junctions. These observations further support that Ca^2+^ release from ER via the PLC-IP3R-CAMK pathway plays a role in disassembly of DE-cad/Arm from adherens junctions. Notably, the results gained with GdCl_3_ treatment indicate that Ca^2+^ entry via Piezo is not required for disassembly or maintenance of DE-cad/Arm at adherens junctions, further supporting that the loss of Arm signals at the lateral side induced by Piezo knockdown (Figure 4) is caused by an impairment of assembly of DE-cad/Arm at adherens junctions.

**Figure 5.**
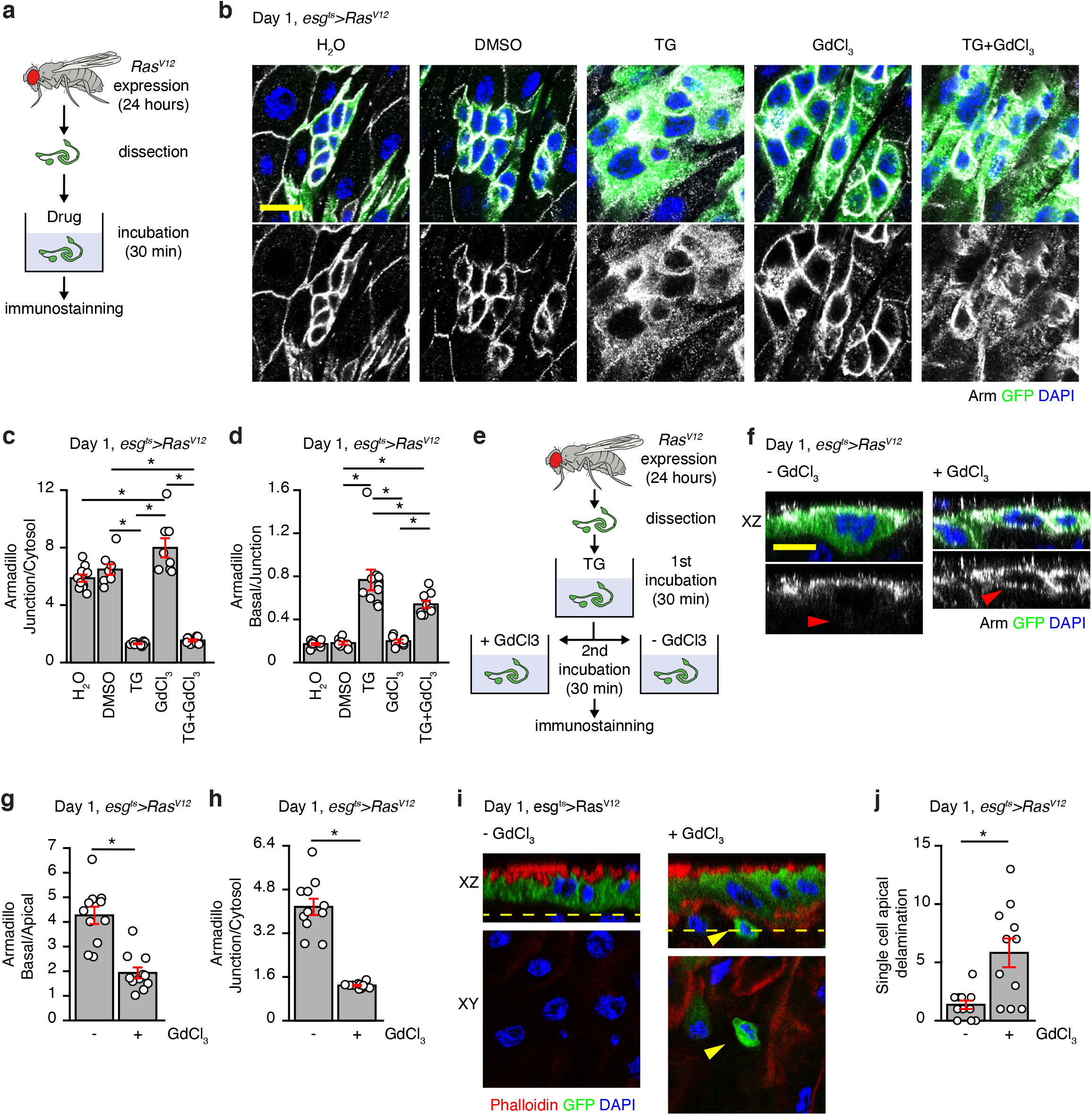
Endoplasmic reticulum calcium release and extracellular calcium entry generate distinct signals regulating the localization of E-cadherin/β-catenin. **a**, Schematic outlining the *ex vivo* midgut manipulation procedure. *esg^ts^>Ras^V12^* midguts are dissected out at day 1 of *Ras^V12^* expression for *ex vivo* treatments for 30 minutes and then stained with anti-Arm antibody. b, Midgut epithelia showing Arm signals (gray). *Ras^V12^* cells are marked by GFP (green), and nuclei are stained with DAPI (blue). Scale bar, 10 μm. c, Ratios of junctional and cytosolic Arm signal intensities. The average fluorescent intensity of Arm at cell-cell junctions are normalized to that in the cytosol. *P*=1.7979×10^-12^ (DMSO - TG), *P*=0.0006 (H_2_O - GdCl_3_), *P*=1.0472×10^-11^ (DMSO - TG+GdCl_3_),\**P*=4.6474×10^-13^ (TG - GdCl_3_), *P*=4.6918×10^-13^ (GdCl_3_ - TG+GdCl_3_). d, Ratios of basal and junctional Arm signal intensities. Average Arm fluorescent intensity in basal membrane was normalized to that in the cell-cell junction. Measurements were acquired from orthogonal views. *P*=7.2906×10^-09^ (DMSO - TG), *P*=2.0376×10^-04^ (DMSO - TG+GdCl_3_), *P=1*.5348×10^-08^ (TG - GdCl_3_), *P*=0.0215 (TG - TG+GdCl_3_), *P*=4.1858×10^-04^ (GdCl_3_ - TG+GdCl_3_). For c and d, each data point represents the mean value of the fluorescent intensities measured from 10 cells in a midgut – a biological replicate. From left to right, *n=*10 (H_2_O), *n*=8 (DMSO), *n*=10 (TG), *n=8* (GdCl_3_), *n*=9 (TG+GdCl3) biological replicates. Mean±SEMs are shown. Statistical analysis was performed using one-way ANOVA with post-hoc Tukey HSD. e, Schematic of the sequential drug treatment experiments. *esg^ts^>Ras^V12^* midguts are dissected out at day 1 of *Ras^V12^* expression for treating TG for 30 minutes. The TG-treated midguts are incubated with or without GdCl3 for 30 minutes. f, Orthogonal views of representative *Ras^V12^* cells treated with TG and then incubated with (right panels) or without GdCl3 (left panels). The basal side of cells is positioned upward, and arrowheads indicate the apical cell surface. Scale bar, 10 μm. g, Ratios of basal and apical Arm signal intensities. *P*=3.0136×10^-05^. h, Ratios of junctional and cytosolic Arm signal intensities. *P*=1.9916×10^-06^. For g and h, each data point represents the mean value of the fluorescent intensities measured from 10 cells in a midgut – a biological replicate. *n*=11 (DMSO), *n*=11 (GdCl3) biological replicates. Mean±SEMs are shown, and \**P*<0.05 (two-tailed unpaired Student’s t-test).i, Orthogonal and cross sections of *esg^ts^>Ras^V12^* midguts. Phalloidin (red) stains actin in the VM, and DAPI (blue) show nuclei. *Ras^V12^* cells are marked with GFP (green). Top panels show orthogonal views (xz), and bottom panels show cross sections (xy) above the apical end of the epithelia. Arrowheads indicate a representative apically delaminating *Ras^V12^* cell. Scale bar, 10 μm. j, Quantification of apical cell delamination. *P*=0.0049. *n*=11(DMSO), *n=*11 (GdCl3) biological replicates. Mean±SEMs are shown. *P<0.05 (two-tailed unpaired Student’s t-test).

### *Ex vivo* gadolinium treatments suggest that Piezo activity is required for assembly of DE-cad/Arm at the basal side and adherens junctions of *Ras^V12^* cells

Interestingly, we observed a marginal, yet significant, decrease in the ratio of basal and junctional Arm signals in *Ras^V12^* cells in TG+GdCl_3_ treatment compared to TG treatment (Figure 5d). Considering that the junctional/cytosol Arm ratios remained comparable between the TG and TG+GdCl_3_ treatments (Figure 5c), the change in the basal/junctional Arm ratio might be due to a decrease in Arm signals at the basal side. This raises an interesting possibility that Piezo might be required for localization of DE-cad/Arm at the basal side after disruption of adherens junctions. Since the presence of TG might cause a continuous disassembly of DE-cad/Arm at adherens junctions and the basal side, we designed a sequential treatment regimen whereby TG was washed out after a 30-minute treatment, and then the midguts were incubated in media either with or without GdCl_3_ for 30 minutes, which would allow *Ras^V12^* cells to recover from the effect of TG, with or without Piezo activity (Figure 5e). Resting TG-treated *Ras^V12^* midguts in media without TG induced strong Arm signals at the basal side of *Ras^V12^* cells (Figure 5f and g), suggesting that resting followed by cytosolic Ca^2+^ increase could induce an assembly of DE-cad/Arm at the basal side. In contrast, resting the TG-treated midguts in media with GdCl_3_ failed to assemble Arm at the basal side (Figure 5f and g), indicating that the mechanosensitive channel activities were required for the basal DE-cad/Arm assembly. Note that the Arm signals at the basal side were not detected as discrete puncta, which might be due to a limitation in recapitulating the native circumstance in our *ex vivo* setting. For instance, considering that DE-cad was not required for formation of invasive protrusions (Figure 2b), assembly of DE-cad/Arm at invasive protrusions might require additional factors, which were not present at day 1 of *Ras^V12^* expression. Additionally, resting the TG-treated midguts in media without GdCl3 increased Arm signals at the lateral side of *Ras^V12^* cells, which was not observed when incubated in media with GdCl3 (Figure 5c and h). Interestingly, incubating the TG-treated midguts in media with GdCl3 significantly increased apical delamination of *Ras^V12^* cells (Figure 5i and j). Previously, it was shown that E-cad was endocytosed from the cell-cell junctions during apical extrusion of *Ras^V12^*-transformed mammalian cells (48). Consistently, apical delamination of *Ras^V12^* cells was predominantly observed at day 2 of *Ras^V12^* expression when adherens junctions were disrupted (27). Thus, further disruption of adherens junctions might account for the increase in apical delamination observed when the TG-treated midguts that were incubated in media with GdCl3 (Figure 5i and j). These results further support that Piezo-mediated calcium signaling is also required for assembly of DE-cad/Arm at adherens junctions. Taken together, our study suggests that the general role of Piezo might be to assemble DE-cad/Arm at the designated compartments depending on the cellular context.

## Discussion

In this study, we describe the unexpected role of DE-cad in the invasiveness of *Ras^V12^* cells during cell dissemination. Without DE-cad, *Ras^V12^* cells couldn’t invade or disseminate *in vivo*. DE-cad/Arm subcellular distribution remodels during dissemination of *Ras^V12^* cells: DE-cad/Arm disassembles at adherens junctions and assembles at invasive protrusions, which are not the conventional locations to detect DE-cad/Arm. Mechanistically, DE-cad controls the function of invasive protrusions, not the formation. Thus, loss of DE-cad results in inability of *Ras^V12^* cells to breach the ECM and the VM. Our findings coincide with the previous clinical reports describing metastasis of *E-cad^+^* cancers (23–26). Padmanaban et al. have demonstrated that E-cad promotes metastasis in multiple models of breast cancer by functioning as a survival factor (15). Our study provides new insight into the molecular mechanism by which E-cad can promote the initial step of metastasis.

Our study elucidates the essential roles of the two intracellular calcium signaling pathways in the invasiveness of *Ras^V12^* cells (Figure S5). The PLC-IP3R-CAMK pathway induces disassembly of DE-cad/Arm at adherens junctions, which recapitulates the functional consequence caused by E-cad loss during epithelial-mesenchymal transition (EMT) (3, 49, 50). Importantly, the Piezo-calpain pathway controls assembly of DE-cad/Arm at invasive protrusions, which is critical for breaching the ECM and the VM. Interestingly, Piezo is required for assembly of DE-cad/Arm at not only invasive protrusions but also the basal side and adherens junctions, suggesting that the general role of calcium entry via Piezo is to control localized assembly of DE-cad/Arm by sensing the mechanical cues in the microenvironment. We have previously reported that Piezo orchestrates dissemination of *Ras^V12^* cells by controlling multiple cellular processes, including invasive protrusion’s function, matrix metalloprotease 1 (MMP1) expression, and bleb-driven movement (27). Additionally, Piezo channels have been shown to be important for matrix degradation in cultured cancer cells (45, 51). A recent preprint suggests that E-cadherin plays a role in invadopodia formation in human cancer cells (52). Thus, the regulation of E-cad assembly by Piezo channels might be a conserved mechanism by which tumor cells coordinate their invasive behavior upon sensing mechanical cues in the microenvironment. Recently, it has been shown that activation of E-cad by injecting an E-cad activating antibody inhibits metastasis in mice presumably by strengthening adherens junctions (16). Our study provides a repertoire of potential targets for strengthening adherens junctions or inhibiting assembly of E-cad at invadopodia to halt the spreading of transformed cells.

## Methods

### *Drosophila* stocks and husbandry

The stocks obtained from the Bloomington *Drosophila* Stock Center (BDSC), the Vienna *Drosophila* Resource Center (VDRC), and the National Institute of Genetics, Japan (NIG) are the following: *UAS-Ras^V12^* (III) (BDSC, 4847), *UAS-DE-cad-i^JF02769^* (BDSC, 27689), *UAS-DE-cad-i^HMS0693^* (BDSC, 32904), *UAS-DE-cad* (BDSC, 655589), *UAS-DE-cad-GFP* (BDSC, 58445), *shg^mTomato^* (referred as *DE-cad^mTomato^;* BDSC, 58789), *UAS-sl-i^HMs00377^* (BDSC, 32385), *UAS-sl-i^HMS00695^* (BDSC, 32906), *UAS-IP3R-i^JF01957^* (BDSC, 25937), *UAS-IP3R-i^GD1676^* (VDRC, 6484), *UAS-IP3R-i^HMC03351^* (BDSC, 51795), *UAS-CaMKII-i* (BDSC, 29401), *UAS-CaMKI-i* (BDSC, 26726), *UAS-piezo-i^8486R-3^* (NIG, 8486R-3), *UAS-CalpA-i^7563R-3^* (NIG, 7563R-3), *UAS-CalpA-i^JF03391^* (BDRC, 29455), *UAS-Actin-mRFP* (BDSC, 24778), and *UAS-Lifeact-mRFP* (BDSC, 58362). We also used *UAS-Ras^V12^*(II) (laboratory stock) and *UAS-mCherry-Moe*.*ABD* (FlyBase, FBtp0108918; gift from Dr. Susan Parkhurst)

We derived transgene expression in intestinal stem cells (ISCs) and enteroblasts (EBs) using *esg-GAL4*, *tub-GAL80^ts^*, *UAS-GFP* (*esg^ts^*) (laboratory stock) as described previously (27, 28). Intestinal epithelial cells manipulated by *esg^ts^* are marked by GFP. Fly crosses were raised in the standard cornmeal-agar medium at 18°C throughout development and adulthood. Three- to ten-day-old non-virgin female flies were used for all the experiments. To induce transgenes, flies were shifted to 29°C for 1-to 2-days prior to dissection.

### Immunostaining

We used the following primary antibodies: mouse anti-Armadillo (1:100; DSHB, N27A1), rat anti-DE-cadherin antibody concentrate (1:50; DSHB, DCAD2), rabbit anti-laminin (1:100; Abcam, ab47650), mouse anti-Mmp1 (1:1000; DSHB, 3B8D12). Alexa Fluor secondary antibodies raised in goat (A11012, A11005, A21244, A21235) were obtained from Thermo Fisher Scientific and used at a dilution of 1:1000. We stained F-actin with phalloidin conjugated to Alexa 594 or 647 (1:1000, Thermo Fisher Scientific, A12381 and A22287). DAPI was used at a 1:2000 dilution (Sigma-Aldrich, D9542).

We fed flies 4% sucrose for approximately 4 hours prior to dissection to remove food from the midgut. Midguts from the adult female flies were then dissected in phosphate buffered saline, pH7.4 (PBS) and fixed for 20 minutes at room temperature in 4% paraformaldehyde (PFA) (Electron Microscopy Sciences, RT15710) diluted in PBS. Midguts were washed three times in PBST (PBS supplemented with 0.2% Triton X-100) and then blocked overnight in PBST supplemented with 5% normal goat serum at 4°C. The tissue samples were then incubated with primary antibody in PBST supplemented with 5% normal goat serum for 2-3 hours at room temperature and rinsed three times in PBST. Secondary antibodies and DAPI were diluted together in PBST supplemented with 5% normal goat serum and incubated with samples at room temperature for 1-2 hours. After secondary incubation, the samples were rinsed three times in PBST and mounted in Vectashield (Vector Laboratories, H1000).

### Image acquisition and quantification

Samples were imaged using a Leica SP8 laser scanning confocal microscope with 40X/1.25 oil or 63X/1.4 oil objective lenses, and confocal stacks (0.8 μm step size or 0.25 μm for higher resolution images) were processed and analyzed using Fiji (ImageJ, National Institutes of Health).

We quantified disseminated cells by counting GFP^+^, DAPI^+^ cells detected on the outer surface of visceral muscle (VM) (27). Confocal stacks (0.8 μm step size) of the posterior midgut were captured in 388 *μ*m x 388 *μ*m fluorescent images. The orthogonal view feature in FIJI was used to determine the position of disseminated cells. Graphical representations of all data points were generated as bar charts with data point overlap in R.

To quantify longitudinal muscle breakage, we visualized the visceral muscles by staining with phalloidin (27). Longitudinal muscle breaks were quantified from one leaflet of the confocal stacks (0.8 μm step size). Fluorescent images (388 μm x 388 μm) were acquired from the posterior midgut with 40X/1.25 oil objective. Graphical representations of all data points were generated as ggplot2 dot plots in R.

Mmp1 fluorescence intensity was quantified from the posterior midgut as described previously (27). Laser settings were kept identical for capturing midgut images. Confocal planes covering one leaflet of the midgut along the apical-basal axis were projected to generate the 388 *μ*m x 388 *μ*m microscope field projection. We collected mean intensity values from three random 100 *μ*m x 100 *μ*m fields per midgut using Fiji and subtracted background values from the area outside surrounding the intestine. Graphical representations of all data points were generated as bar charts with data point overlap in R.

To measure fluorescent intensity of Laminin B1, we generated z-projection (388 *μ*m x 388 *μ*m) of confocal stacks covering one leaflet of the posterior midgut. Mean gray values from three random 100 *μ*m x 100 *μ*m regions per midgut were collected. The background was then subtracted using values from the area outside the intestine. Graphical representations of all data points were generated as bar charts with data point overlap in R.

We defined tumors as epithelial layers having over 80% of the layer comprising of GFP-positive cells. Images were acquired from the R5 region of the posterior midgut with the 40x/1.25 oil objective covering 388 *μ*m x 388 *μ*m in area. Graphical representations of all data points were generated in the percentage stacked bar chart in R.

### *Ex vivo* drug treatment and quantification

For the *ex vivo* experiments, we essentially used *Drosophila* Adult Hemolymph-like saline (AHLS) comprising 2mM CaCl_2_, 5mM KCl, 5mM HEPES, 8.2mM MgCl_2_,108mM NaCl, 4mM NaHCO_3_, 1mM NaH_2_PO_4_, 10mM Sucrose, and 5mM Trehalose (53). The solution was adjusted to pH 7.5 and stored at 4C. We dissolved Thapsigargin (TG) (Sigma-Aldrich, T9033) in DMSO and GdCl_3_ (EMD Millipore, G75325G) in water to make stock solutions.

Ras^V12^ was expressed with *esg^ts^* for 24 hours and dissected in AHLS to prepare the whole midguts. The dissected midguts were then incubated in AHLS supplemented with 4% Fetal Bovine Serum (Life Technologies, 16140071) for drug treatment for 30 minutes at room temperature in unsealed glass dissection wells. Intestines were then washed in PBS, fixed in 4% PFA, and immunostained. Fluorescent images (388 μm x 388 μm) were captured from the posterior midguts. Laser settings were kept identical for all conditions. For sequential treatment experiments, dissected midguts were incubated with 1μM TG for 30 minutes and then washed three times in AHLS with 4% FBS. The TG-treated midguts were then incubated in AHLS with 4% FBS containing either 100μM GdCl3 or solvent for 30 minutes. Midguts were washed in PBS and followed by immunostaining. All steps occurred in unsealed wells at room temperature.

To assess junction integrity, we measured Armadillo (Arm) signal intensity from 10 cells per midgut. Arm signal intensity was measured from a representative cross-section for a given cell using the plot profile feature in FIJI. We measured Arm signal intently along a line drawn from the cell boundary to the nuclear boundary. We took the measured value at the midpoint of the line as the cytosolic Arm signal intensity and compared to the value obtained at the cell boundary/junction. For statistical analysis, one-way ANOVA with post-hoc Tukey HSD (Honestly Significant Difference) was performed using R.

To assess the enrichment of Arm at the basal surface relative to junction, we used a representative orthogonal view of a given cell. We measured Arm signal intensity at the basal boundary and compared to Arm signal intensity measured at the lateral boundary of the cell. The ratios of Basal and Junctional signal intensities were presented, and one-way ANOVA with post-hoc Tukey HSD (Honestly Significant Difference) was performed using R.

To assess the Arm signal ratio between basal and apical surfaces, Arm signals were quantified from ten cells per midgut. We measured signal intensity values from a representative orthogonal view of given cell by tracing cell’s basal and apical boundaries. The ratios of basal and apical Arm signals were calculated, and two-tailed unpaired t-test was performed for statistical analysis.

To quantify apically delaminating cells, we used the orthogonal view in FIJI to determine the midgut epithelial boundary-outlined with phalloidin staining. GFP^+^, DAPI^+^ cells detected outside the midgut epithelial boundary were counted. For statistical analyses, two-tailed unpaired t-tests were performed.

### Statistics and reproducibility

Statistical differences between groups of data were analyzed using a series of two-tailed unpaired Student t-test or one-way ANOVA with post-hoc Tukey HSD (Honestly Significant Difference). All statistical analyses and data graphics were done in ‘R’ software (version 1.3.1093). All quantified experiments represent at least three biologically independent samples. Level of significance are depicted by asterisks in the figures: \**P*<0.05. *P* values are indicated in the figure legends. Sample sizes were chosen empirically based on the observed effects and listed in the figure legends. All quantifications are from the posterior midguts of the adult female flies.

## Data availability

The data supporting the finding of this study are available within the paper and its Supplementary Information. Source data are provided with this paper.

## Acknowledgements

We thank Dr. Susan Parkhurst for sharing fly stocks, Dr. Jiae Lee for helping with imaging, and Eric So for the technical assistance. This work is supported by NIH R35GM128752 to Y.V.K. and by NIH T32CA080416 to A.J.H.C.

## Author Contributions

A.J.H.C., Y.V.K., and G.M.G. designed experiments, analyzed data and wrote manuscript. A.J.H.C. performed experiments.

