## Supplementary Figures 1-5 for "The roles of distinct Ca^2+^ signaling mediated by Piezo and inositol triphosphate receptor (IP3R) in the remodeling of E-cadherin during cell dissemination"

**Figure S1. DE-cadherin is dispensable for delamination of *Ras$^{V12}$* cells. a,** Posterior midguts at day 2 of transgene expression by *esg$^{ts}$*. Two DE-cad RNAi lines used to knockdown DE-cad are indicated as *DE-cad-i$^{JF02769}$* or *DE-cad-i$^{HMS0693}$*. Cells manipulated by *esg$^{ts}$* are labeled with GFP (green), and nuclei are stained with DAPI (blue). Scale bar, 50 $\mu$m. **b,** Representative images of the posterior midgut at day 2 of transgene expression. Cells manipulated with *esg$^{ts}$* express GFP (green). VM is stained with phalloidin (red) and nuclei with DAPI (blue). Images are single confocal stacks showing the longitudinal section of the midgut. Representative delaminating GFP$^{+}$ cells are indicated with arrowheads. Scale bar, 50 $\mu$m.

**Figure S2. DE-cadherin/Armadillo localizes at invasive protrusions.**

**a,** Basal and orthogonal views showing DE-cad signals (gray). Day 1 $esg^{ts}$>+ (control) and $esg^{ts}$>$Ras^{V12}$ midguts were stained with anti-DE-cad antibody. Arrowheads indicate DE-cad signals at cell-cell junctions. Asterisks indicate nonspecific signals in the visceral muscle layer. Scale bar, 5 μm. **b,** Basal and orthogonal views showing Arm signals (gray). Day 1 $esg^{ts}$>+ (control) and $esg^{ts}$>$Ras^{V12}$ midguts were stained with anti-Arm antibody. Arrowheads indicate Arm signals at cell-cell junctions. Scale bar, 5 μm. **c**, Basal and orthogonal views showing DE-cad signals (red) at day 2 of transgene expression. yellow arrowheads indicate Arm signals at cell-cell junctions. White arrowheads show DE-cad signal puncta at the basal side of $Ras^{V12}$ cell. Scale bar, 10 μm. **d**, DE-cad (red) and Arm (gray) co-localization at invasive protrusions. DE-cad signals (red) was detected by using a DE-cad protein trap line ($DE\text{-}cad^{mTomato}$), which produces DE-cad-mTomato under control of the native *DE-cad* regulatory sequence. Arm signals (gray) were detected by staining with anti-Arm antibody. Arrowheads indicate basal puncta showing both DE-cad-mTomato and Arm signals. Scale bar, 10 μm. In all the orthogonal sections, the basal side of the epithelia is position upward.

**Figure S3. Depletion of the PLC-IP3R-CAMK pathway components induces midgut tumors.**

**a**, Percentage stacked bar chart of midguts with epithelial tumors. *n*=10 ($esg^{ts}$), *n*=14 ($esg^{ts}$>$Ras^{V12}$), *n*=14 ($esg^{ts}$>$Ras^{V12}$, $sl\text{-}i^{HMS00377}$), *n*=19 ($esg^{ts}$>$Ras^{V12}$, $sl\text{-}i^{HMS00695}$), *n*=8 ($esg^{ts}$>$Ras^{V12}$, $IP3R\text{-}i^{JF01957}$), *n*=14 ($esg^{ts}$>$Ras^{V12}$, $IP3R\text{-}i^{GD1676}$), *n*=9 ($esg^{ts}$>$Ras^{V12}$, $IP3R\text{-}i^{HMC03351}$), *n*=10 ($esg^{ts}$>$Ras^{V12}$, *CaMKI-i, CaMKII-i*) biological replicates. **b**, Representative midgut longitudinal sections. Transgenes were induced for 2 days with $esg^{ts}$. Cells manipulated with $esg^{ts}$ express GFP (green). Visceral muscles are visualized with Phalloidin (red), and nuclei are stained with DAPI (blue). Arrowheads indicate representative delaminating $GFP^{+}$ cells. Scale bar, 50 $\mu$m. **c**, Percentage stacked bar chart of midguts with epithelial tumors. *n*=10 ($esg^{ts}$), *n*=14 ($esg^{ts}$>$Ras^{V12}$), *n*=14 ($esg^{ts}$>$Ras^{V12}$, $sl\text{-}i^{HMS00377}$), *n*=19 ($esg^{ts}$>$Ras^{V12}$, $sl\text{-}i^{HMS00695}$), *n*=15 ($esg^{ts}$>$Ras^{V12}$, $norpA\text{-}i^{JF01518}$), *n*=16 ($esg^{ts}$>$Ras^{V12}$, $norpA\text{-}i^{JF01713}$), *n*=9 ($esg^{ts}$>$Ras^{V12}$, $Plc21c\text{-}i^{HMS00436}$), *n*=17 ($esg^{ts}$>$Ras^{V12}$, $Plc21c\text{-}i^{JF01210}$), *n*=10 ($esg^{ts}$>$Ras^{V12}$, $Plc21c\text{-}i^{HMS00600}$) biological replicates.

**Figure S4. Basal Arm puncta are occasionally observed in *Ras$^{V12}$*, *CaMKI-i*, *CaMKII-i* cells.**

Arm signals are in gray. Red arrowheads indicate representative basal Arm puncta, and yellow arrowheads show cell-cell junctions. DAPI is in blue. Transgenes are expressed for 2 days with *esg$^{ts}$*. Scale bar 10 $\mu$m. The basal side of the epithelia is position upward in the orthogonal section.

**Figure S5. E-cadherin/β-catenin regulation by intracellular calcium signaling in *Ras$^{V12}$* cells**

E-cadherin/β-catenin localizes to adherens junctions in control epithelial cells. During dissemination of *Ras$^{V12}$* cells, E-cadherin/β-catenin disassembles at adherens junctions and assembles at invasive protrusions. The disassembly of E-cadherin/β-catenin at adherens junctions is induced by intracellular calcium signaling caused by ER $Ca^{2+}$ release via the PLC-IP3R pathway. The assembly of E-cadherin/β-catenin at invasive protrusions is induced by $Ca^{2+}$ entry via Piezo. Given the role of Piezo in mechanosensation, Piezo-mediated assembly of E-cadherin/β-catenin may allow *Ras$^{V12}$* cells to adjust their invasive behavior by sensing mechanical cues in their microenvironment.

### Figure S1

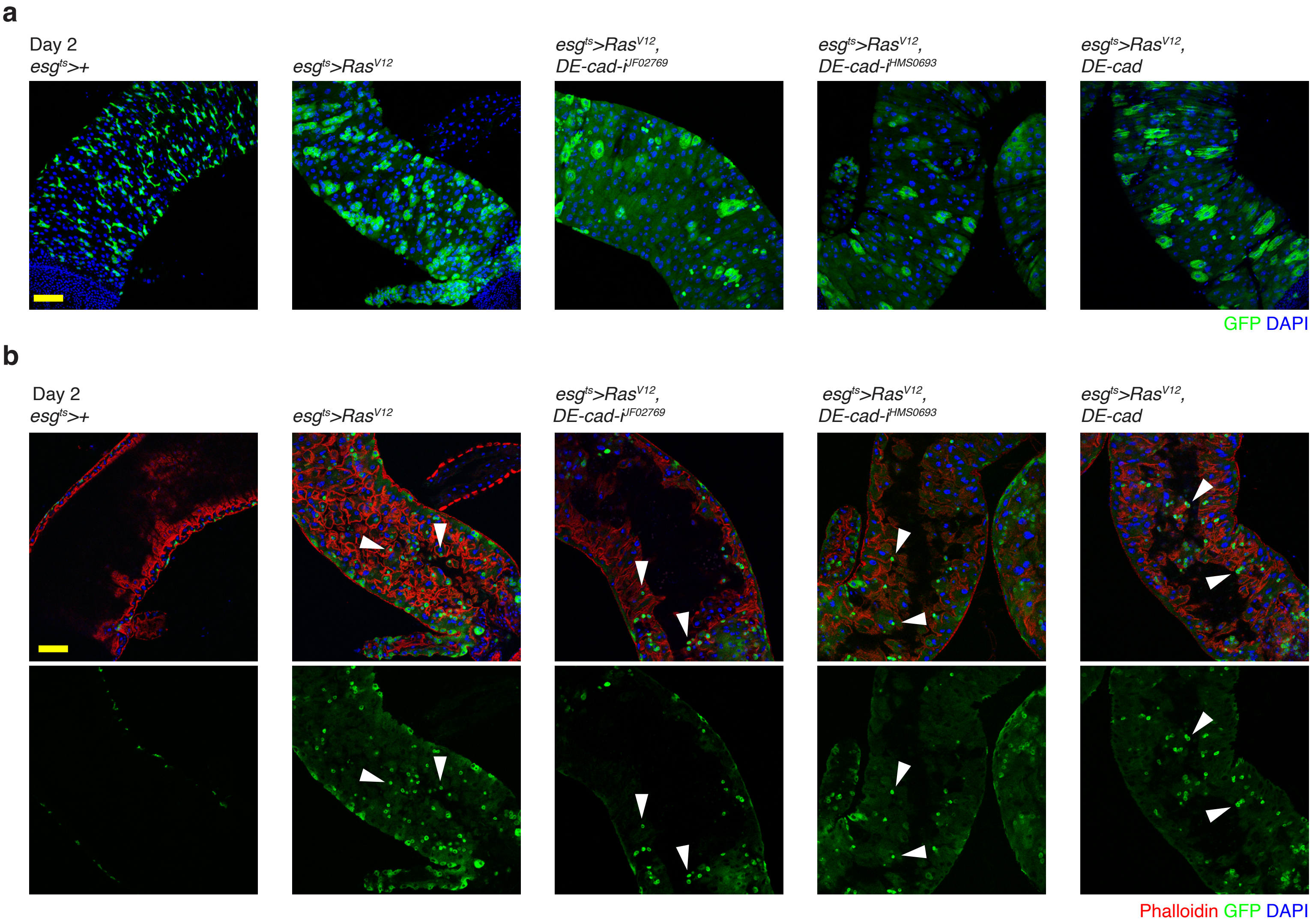

a
Day 2
esg^ts>+
esg^ts>Ras^V12
esg^ts>Ras^V12, DE-cad-i^JF02769
esg^ts>Ras^V12, DE-cad-i^HMS0693
esg^ts>Ras^V12, DE-cad
GFP DAPI
b
Day 2
esg^ts>+
esg^ts>Ras^V12
esg^ts>Ras^V12, DE-cad-i^JF02769
esg^ts>Ras^V12, DE-cad-i^HMS0693
esg^ts>Ras^V12, DE-cad
Phalloidin GFP DAPI

### Figure S2

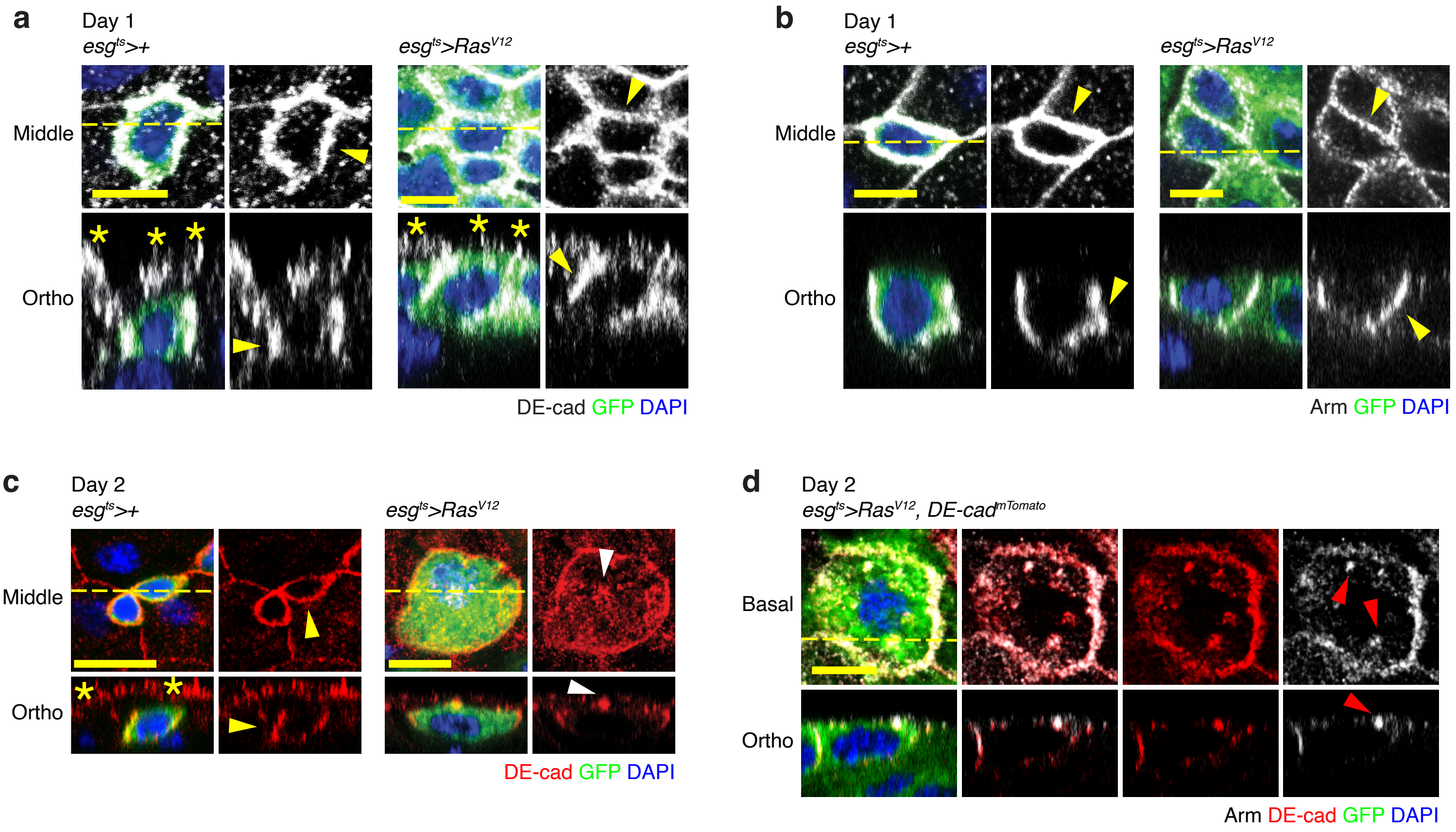

a
Day 1
esg^ts>+
esg^ts>Ras^V12
Middle
Ortho
DE-cad GFP DAPI
b
Day 1
esg^ts>+
esg^ts>Ras^V12
Middle
Ortho
Arm GFP DAPI
c
Day 2
esg^ts>+
esg^ts>Ras^V12
Middle
Ortho
DE-cad GFP DAPI
d
Day 2
esg^ts>Ras^V12, DE-cad^mTomato
Basal
Ortho
Arm DE-cad GFP DAPI

### Figure S3

**a**

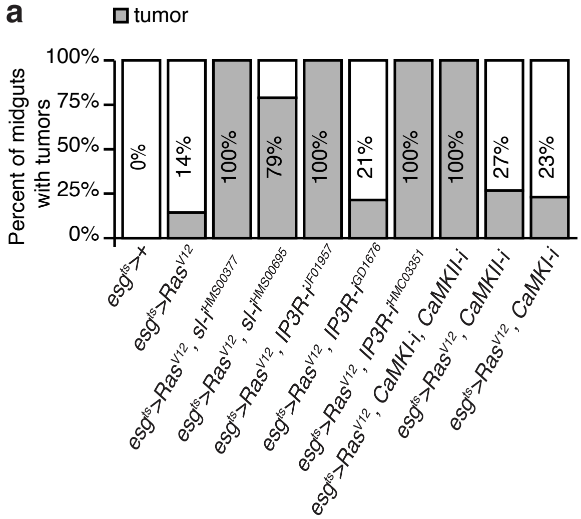


**b**

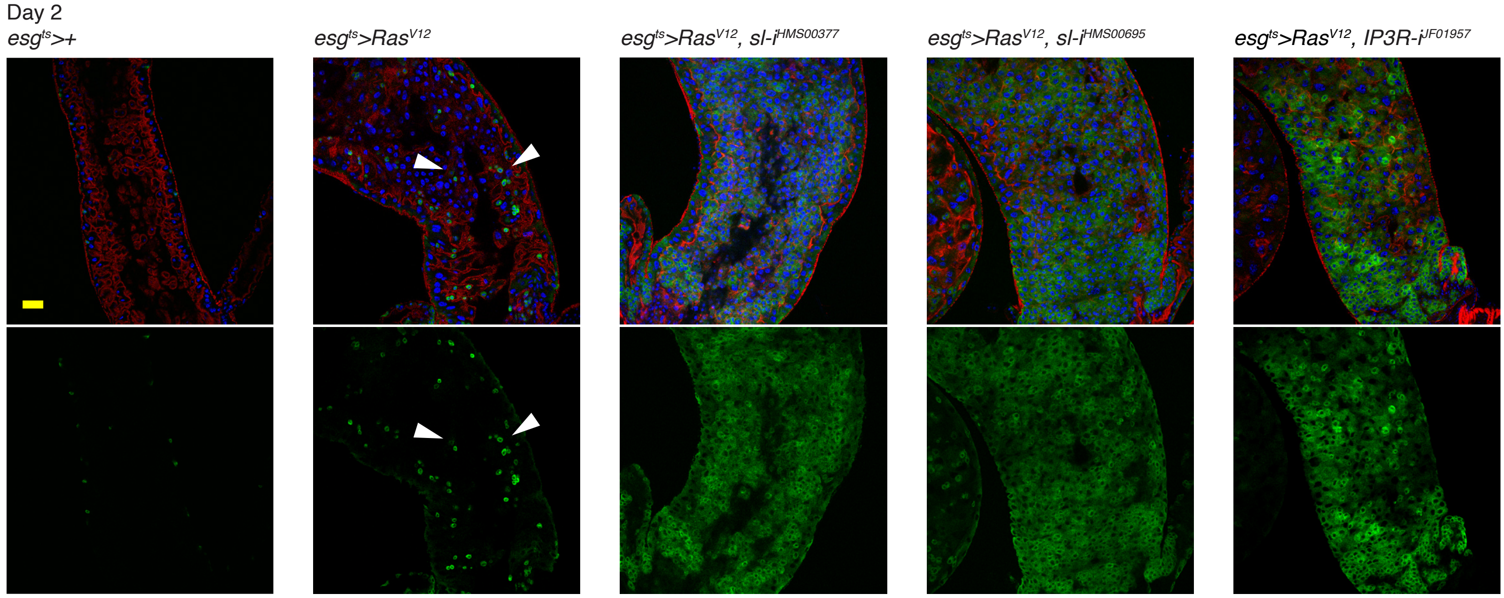


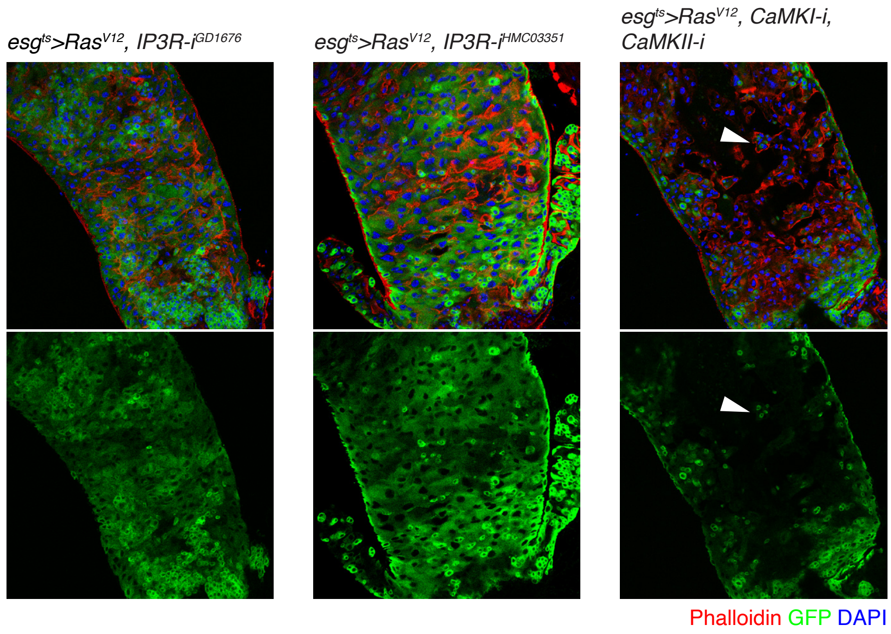


**c**

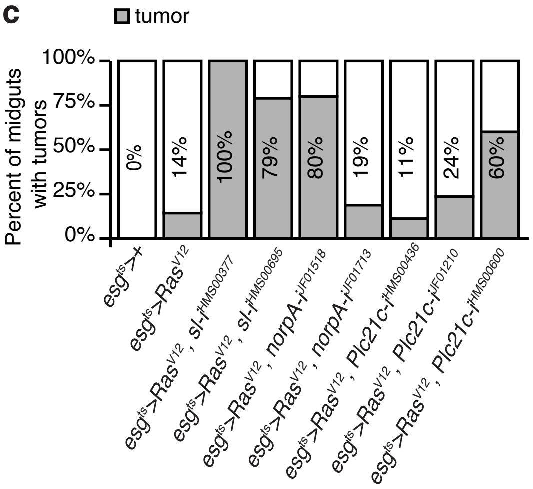

### Figure S4

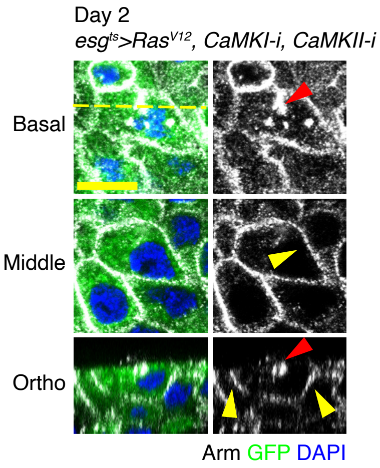

Day 2
*esg^ts>Ras^V12, CaMKI-i, CaMKII-i*
Basal
Middle
Ortho
Arm GFP DAPI

### Figure S5

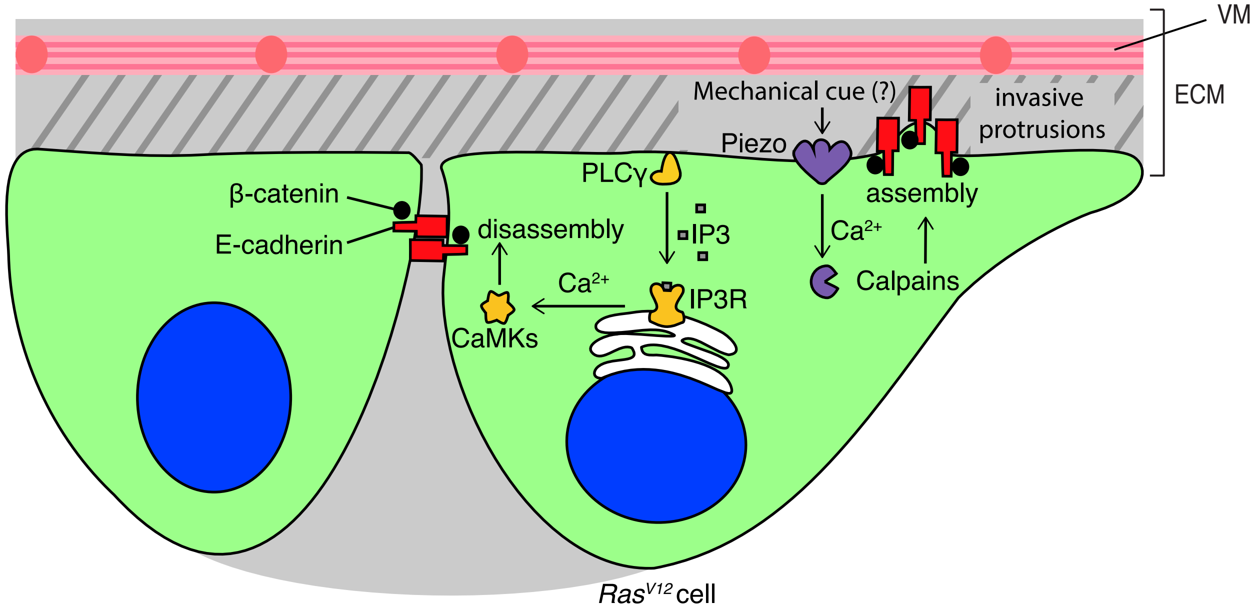
VM
ECM
Mechanical cue (?)
invasive
protrusions
Piezo
PLCγ
β-catenin
E-cadherin
disassembly
assembly
IP3
Ca2+
Ca2+
Calpains
IP3R
CaMKs
RasV12 cell
